# A note on detecting statistical outliers in psychophysical data

**DOI:** 10.1101/074591

**Authors:** Pete R. Jones

## Abstract

This paper considers how best to identify statistical outliers in psychophysical datasets, where the underlying sampling distributions are unknown. Eight methods are described, and each is evaluated using Monte Carlo simulations of a typical psychophysical experiment. The best method is shown to be one based on a measure of absolute-deviation known as *S*_*n*_. This method is shown to be more accurate than popular heuristics based on standard deviations from the mean, and more robust than non-parametric methods based on interquartile range. Matlab code for computing *S*_*n*_ is included.

## 1. The problem of outliers

A statistical outlier is an observation that diverges abnormally from the overall pattern of data. They are often generated by a process qualitatively distinct from the main body of data. For example, in psychophysics, spurious data can be caused by technical error, faulty transcription, or — perhaps most commonly — participants being unable or unwilling to perform the task in the manner intended (e.g., due to boredom, fatigue, poor instruction, or malingering). Whatever the cause, statistical outliers can profoundly affect the results of an experiment^1^, making similar populations appear distinct (Fig 1A, *top panel*), or distinct populations appear similar (Fig 1A, *bottom panel*). For example, it is tempting to wonder how many ‘developmental’ differences between children and adults are due to a small subset of non-compliant children.

**Fig 1.**
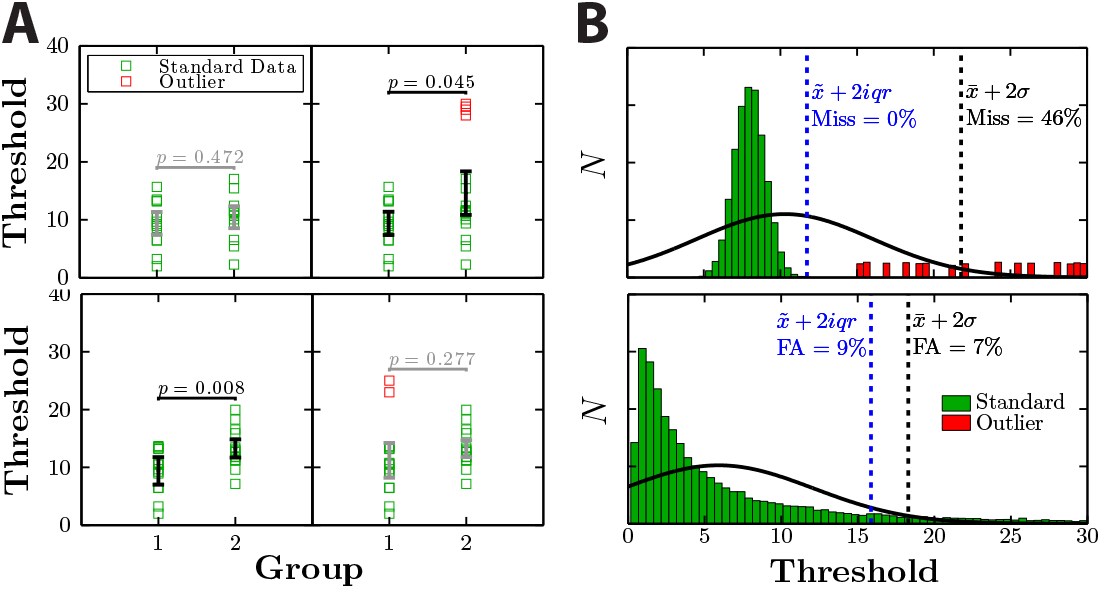
Examples of (***A***) how the presence outliers can qualitatively affect the overall pattern of results, and (***B***) common errors made by existing methods of outlier identification heuristics. *P*-values pertain to the results of between-subject *t*-tests. See body text for details.

## 2. General approaches and outstanding questions

One way to militate against outliers is to only ever use non-parametric statistics, which have a high breakdown point^2^, and so tend to be robust against extreme values. In reality though, non-parametric methods are often impractical, since they are less powerful, less well understood, and less widely available than their parametric counterparts. Alternatively then, many experimenters identify and remove outliers ‘manually’, using some unspecified process of ‘inspection’. This approach is not without merit. However, when used in isolation, manual inspection is susceptible to bias and human error, and it precludes rigorous replication or review. Finally then, statistical outliers can be identified numerically. If the underlying sampling distribution is known, then it is trivial to set a cutoff based on the likelihood of observing a given data point. However, when the sampling distribution is unknown, researchers are often compelled to use numerical heuristics, such as “was the data point more than *N* standard deviations from the mean?”. Currently, a plethora of such heuristics exist in common usage. It is unclear which method works best, and at present unscrupulous individuals are free to pick-and-choose whichever yields the outcome they expect/desire. The goal of this work was therefore (*i*) to describe what methods are currently available for identifying statistical outliers (in datasets generated from an unknown sampling distribution), and (*ii*) to use simulations to assess how well each method performs in a typical psychophysical context.

## 3. State-of-the-art methods for identifying statistical outliers

Here we describe eight methods for identifying statistical outliers. Five of these methods are also shown graphically in Fig 3.

***SD*** *x*_*i*_=outlier if it lies more than *λ* standard deviations, *σ*, from the mean, 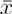:

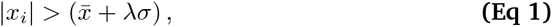

where *λ* is typically between 2 (liberal) and 3 (conservative). This is one of the most commonly used heuristics, but is theoretically flawed. Both the 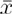 and *σ* terms are easily distorted by extreme values, meaning that more distant outliers may ‘mask’ lesser ones. This can lead to false negatives (identifying outliers as genuine data; Fig 1B, *top panel*). The method also assumes symmetry (i.e., attributes equal importance to positive and negative deviations from the center), whereas psychometric data are often skewed. This can lead to false positives (identifying genuine data as outliers; Fig 1B, *bottom panel*). Furthermore, while *SD* does not explicitly require normality, the ±*λσ* bracket may include more or less data than expected if the data are not Gaussian distributed. For example, ±2*σ* includes 95% of data when Gaussian distributed, but as little as 75% otherwise (*Chebyshev’s inequality*).

***GMM*** *x*_*i*_ = outlier if it lies more than *λ* standard deviations from the mean *of the primary component of a Gaussian Mixture Model:*

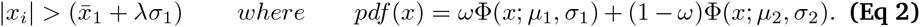

An obvious extension to *SD:* The two methods are identical, except that when fitting the parameters to the data, the *GMM* model also includes a secondary component designed to capture any outliers (see Fig 3). The secondary component is not used to identify outliers per se, but prevents extreme values from distorting the parameters of the primary component. In practice the fit of the secondary component must be constrained to prevent it ‘absorbing’ non-outlying points.

***rSD*** Same as *SD*, but applied recursively until no additional outliers are identified:

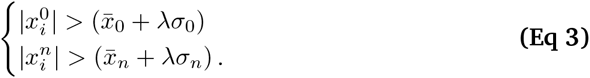

This approach aims to solve the problem of masking by progressively peeling away the most extreme outliers. However, like *SD*, it remains intolerant to non-Gaussian distributions. In situations where samples are sparse/skewed, this approach therefore risks aggressively rejecting large quantities of genuine data (see Fig 1B). Users typically attempt to compensate for this by using a relatively high criterion level, and/or by limiting the number of recursions (e.g., *λ* ≥ 3, *n*_max_=3).

***IQR*** *x*_*i*_=outlier if it lies more than *λ* times the interquartile range from the median:

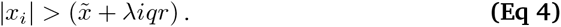

This is a non-parametric analog of the *SD* rule: substituting median and *iqr* for mean and standard deviation. Unlike *SD*, the key statistics are relatively robust. Thus, the breakdown points for 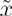 and *iqr* are 50% and 25% (respectively), meaning that outliers can constitute up to 25% of the data before the statistics start to be distorted^3^. However, like *SD*, the *IQR* method only considers absolute deviation from the center. It is therefore insensitive to any asymmetry in the sampling distribution (Fig 1B, *bottom*).

***prctile*** *x*_*i*_ =outlier if it lies above the *λ*^th^ percentile, or below the (1 – *λ*)^th^:

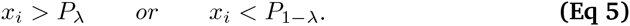

This effectively ‘trims’ the data, rejecting the most extreme points, irrespective of their values. Unlike *IQR*, this method is sensitive to asymmetry in the sampling distribution. But it is otherwise crude in that it ignores any information contained in the spread of the data points. The *prctile* method also begs the question in that the experimenter must estimate, *a priori*, the number of outliers that will be observed. If *λ* is set incorrectly, genuine data will be excluded, or outliers missed.

***Tukey*** *x*_*i*_=outlier if it lies more than *λ* times the iqr from the 25^th^/75^th^ percentile:

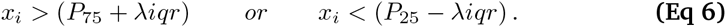

Popularized by John W. Tukey, this attempts to combine the best features of the *IQR* and *prctile* method. The information contained in the spread of data, *iqr*, is combined with the use of lower/upper quartile ‘fences’ that provide some sensitivity to asymmetry.

***MAD*_*n*_** *x*_*i*_=outlier if it lies farther from the median than λ times the median absolute distance [MAD] of every point from the median:

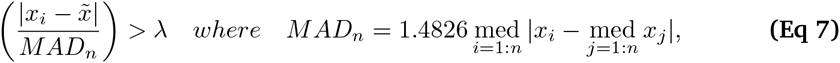

where 1.4826 is simply a scaling factor, used for consistency with the standard deviation over a Gaussian distribution (see Ref [3]). Unlike the non-parametric methods described previously, this method uses MAD rather than *iqr* as the measure of spread. This makes this method more robust, as the MAD statistic has the best possible breakdown point (50%, versus 25% for *iqr*). However, as with *IQR, MAD*_*n*_ assumes symmetry, only considering the absolute deviation of datapoints from the center.

***S***_*n*_ *x*_*i*_=outlier if the median distance of *x*_*i*_ from all other points, is greater than *λ* times the median distance of every point from every other point:

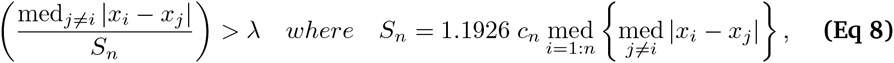

where 1.1926 is again for consistency with the standard deviation, and *c*_*n*_ is a finite population correction parameter (see Ref [3]). Like MAD, the *S*_*n*_ term is maximally robust. However, this method differs from *MAD*_*n*_ in that *S*_*n*_ considers the typical distance between all data points, rather than measuring how far each point is from a central value. It therefore remains valid even if the sampling distribution is asymmetric. The historic difficulty with *S*_*n*_ is its computational complexity [O(*n*^2^)]. However, for most psychophysical applications processing times are on the order of milliseconds, given modern computing power.

## 4. Comparison of techniques using simulated psychophysical observers

To assess the eight methods described in *Section 3*, each was applied to random samples of data prelabeled as either ‘bad’ (should be excluded) or ‘good’ (should not be excluded). Rather than simply specifying arbitrary sampling distributions for these categories, we generated data by simulating a typical two-alternative forced-choice [2AFC] experiment in which a 2-down 1-up transformed staircase^4^ was applied to *N* simulated observers.

Each observer consisted of a randomly generated psychometric function (Fig 2), which was used to make stochastic, trial-by-trial responses based on the current stimulus level and a random sample of additive internal noise. Trial-by-trial response data were then processed and analyzed as if from a human participant. Of the *N* observers, *X%* were ‘non-compliant’ (on average, their psychometric functions had a higher mean, variance, and lapse-rate), and were thus likely to produce outlying data points (i.e., estimates of 70.7% threshold: Fig 3, *red bars*). The remaining observers were ‘compliant’ (on average lower mean, variance, and lapse-rate), and produced the main distribution of ‘good’ data (Fig 3, *green bars*).

**Fig 2.**
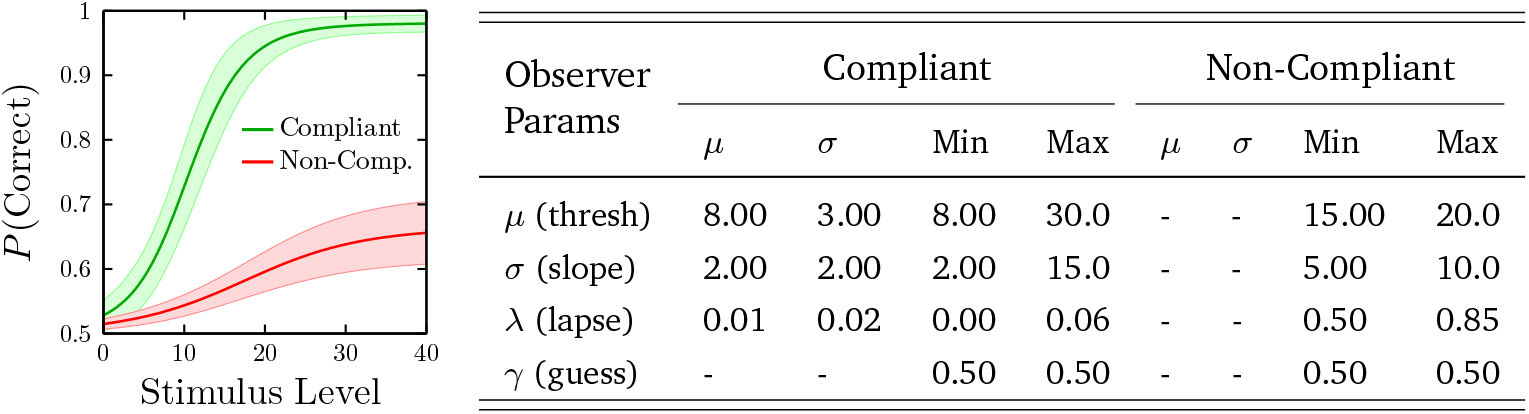
Mean [±1 SD] empirical cumulative distributions for simulated observers. For all individuals, the shape of the underlying psychometric function was logistic. Guess rate was fixed at 50%. Other parameters were independently randomly generated for each simulated observer, using either a truncated Gaussian (compliant) or uniform distribution (non-compliant) — see table for parameters.

**Fig 3.**
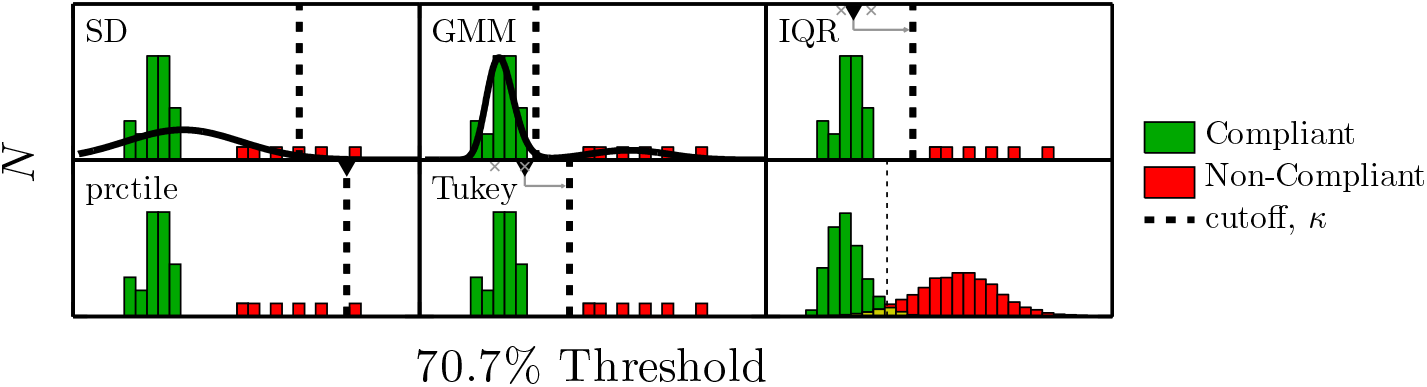
Methods. Each observer’s trial-by-trial responses were used to generate an estimate of their 70.7% correct threshold. *X*% of threshold estimates came from ‘non-compliant’ simulated observers (here: *N*=32, *X*%=19). Each of eight methods in *Section 3* were then used to identify which observations were generated by ‘non-compliant’ observers (i.e., likely statistical outliers). Only five methods are depicted here, as the other three (*rSD*, *MAD*_*n*_ and *S*_*n*_) have no obvious graphical analog. The final panel shows the full sampling distributions over 20,000 trials, and the ideal unbiased classifier, for which: Hit rate=0.97, False Alarm=0.05.

The number of observers, *N*, took the values 〈8, 32, 128〉, representing small, medium, and large sample size. The number of non-compliant observers varied from 0 to 50% of *N* (e.g., 〈0,1,…, 16〉, when when *N=32*). For each condition, 2,000 independent simulations were run, for a total of 108K simulations.

### Results and Discussion

The results are shown in Fig 4. We begin by considering only the case where *N=32* (Fig 4, *middle column*), before considering the effect of sample size.

**Fig 4.**
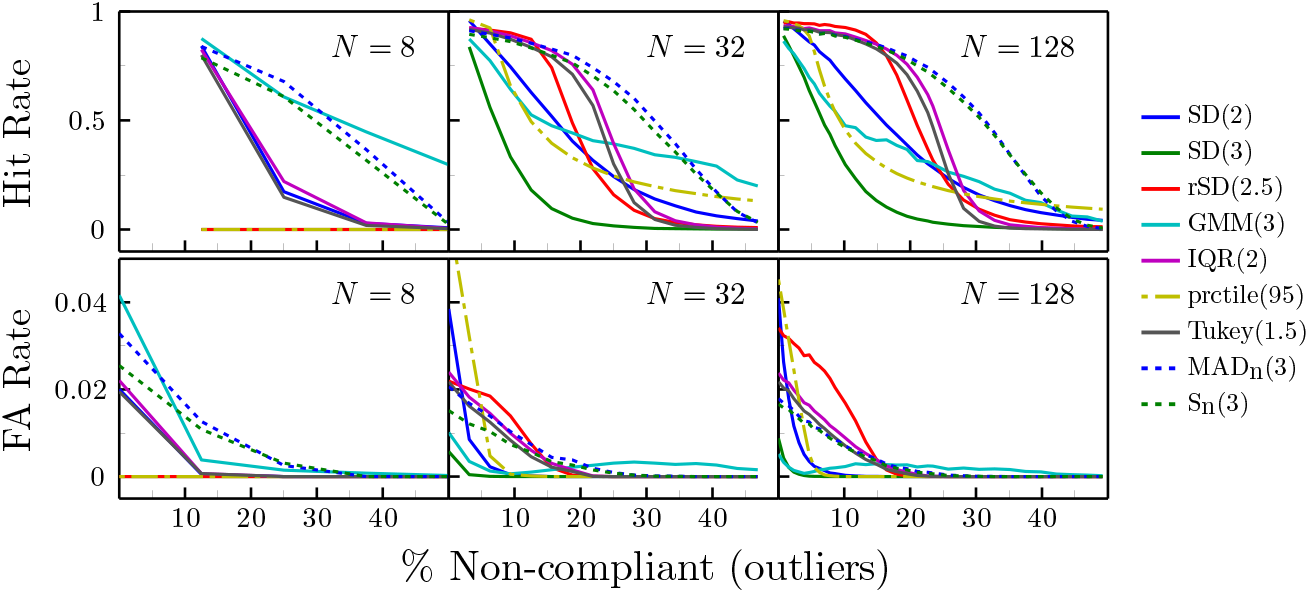
Simulation results. The eight classifiers described in *Section 3* were used to distinguish between random samples of ‘compliant’ and ‘non-compliant’ simulated observers (see Fig 3). Numbers in parentheses indicate the criterion level, *λ*, used by each classifier.

As expected, the *SD* rule proved poor. When *λ*=3, it was excessively conservative –seldom exhibiting false alarms, but missing the great majority of outliers, particularly as the number of outliers increased. Lowering the criterion to *λ*=2 yielded more reasonable results. However, *SD* still exhibited a lower hit rate than most other methods, and also exhibited a high false alarm rate when there were few/no outliers. The modified *GMM* and *rSD* rules exhibited increased robustness and accuracy, respectively. However, compared to non-parametric methods, they were generally only more sensitive than the *prctile* method, which was only accurate when the predefined exclusion rate matched the true number of outliers exactly.

The two *iqr*-based methods, *IQR* and *Tukey*, exhibited high sensitivity when the number of outliers was low (≤20%). However, as expected, they exhibited a marked deterioration in hit rates when the number of outliers increased beyond 20% (i.e., in accordance with the 25% breakdown point for *iqr*).

The two median-absolute-deviation-based methods, *MAD*_*n*_ and *S*_*n*_, were as sensitive as all other methods when outliers were few (≤20%), and were more robust than the *iqr* methods – continuing to exhibit high hit rates and few false alarms even when faced with large numbers of outliers. Compared to each other, *MAD*_*n*_ and *S*_*n*_ performed similarly. However, the S_**n**_ statistic makes no assumption of symmetry, and so ought to be superior in situations where the sampling distribution is heavily skewed.

We turn now to how sample size affected performance. With large samples (*N*=128), the pattern was largely unchanged from the medium sample-size case (*N*=32), except that *rSD* exhibited a marked increase in false alarms, making it an unappealing option. With small samples (*N=8*), the *prctile* and *rSD* methods became uniformly inoperable, while most other methods were unable to identify more than a single outlier. The *MAD*_*n*_ and *S*_*n*_ methods, however, remained relatively robust, and generally performed well, though they did exhibit an elevated false alarm rate when there were few/no outliers. It may be that this could be rectified by increasing the criterion, *λ*, as a function of *N*, however this was not investigated here. The *GMM* method also performed well overall in the small-sample condition. However, it did also exhibit the highest false alarm rate when there were no outliers, and was only more sensitive than *MAD*_*n*_ or *S*_*n*_ when the proportion of outliers was extremely high (>33%).

## 5. Summary and concluding remarks

Of the eight methods considered, *S*_***n***_ proved the most sensitive and robust. Specific situations were observed in which other heuristics performed as-well-as or even better than *S*_***n***_: for example, when the sample size was large (*rSD*), or when the proportion of outliers was very low (*IQR, Tukey*) or very high (*GMM*). However, most methods were less sensitive in than *S*_*n*_ in the majority circumstances, and failed precipitously in some circumstances, making them unattractive alternatives. The related method *MAD*_*n*_ also proved strong, and can be considered a good method for identifying outliers, as has been noted previously by others^5^. However, as discussed in *Section 3, MAD*_*n*_ assumes a symmetric sampling distribution, and so would not be expected to perform as well in situations where the sampling distribution is very heavily skewed (e.g., when dealing with reaction time data). The popular *SD* metric proved particularly poor in all circumstances, and should never be used. In short, *S*_*n*_ appears to provide the best means of identifying statistical outliers when the underlying sampling distribution is unknown. Its use may be particularly beneficial for researchers working with small/irregular populations such as children, animals, or clinical cohorts. MATLAB code for computing *S*_***n***_ is provided in Listing 1.

**Listing 1.**
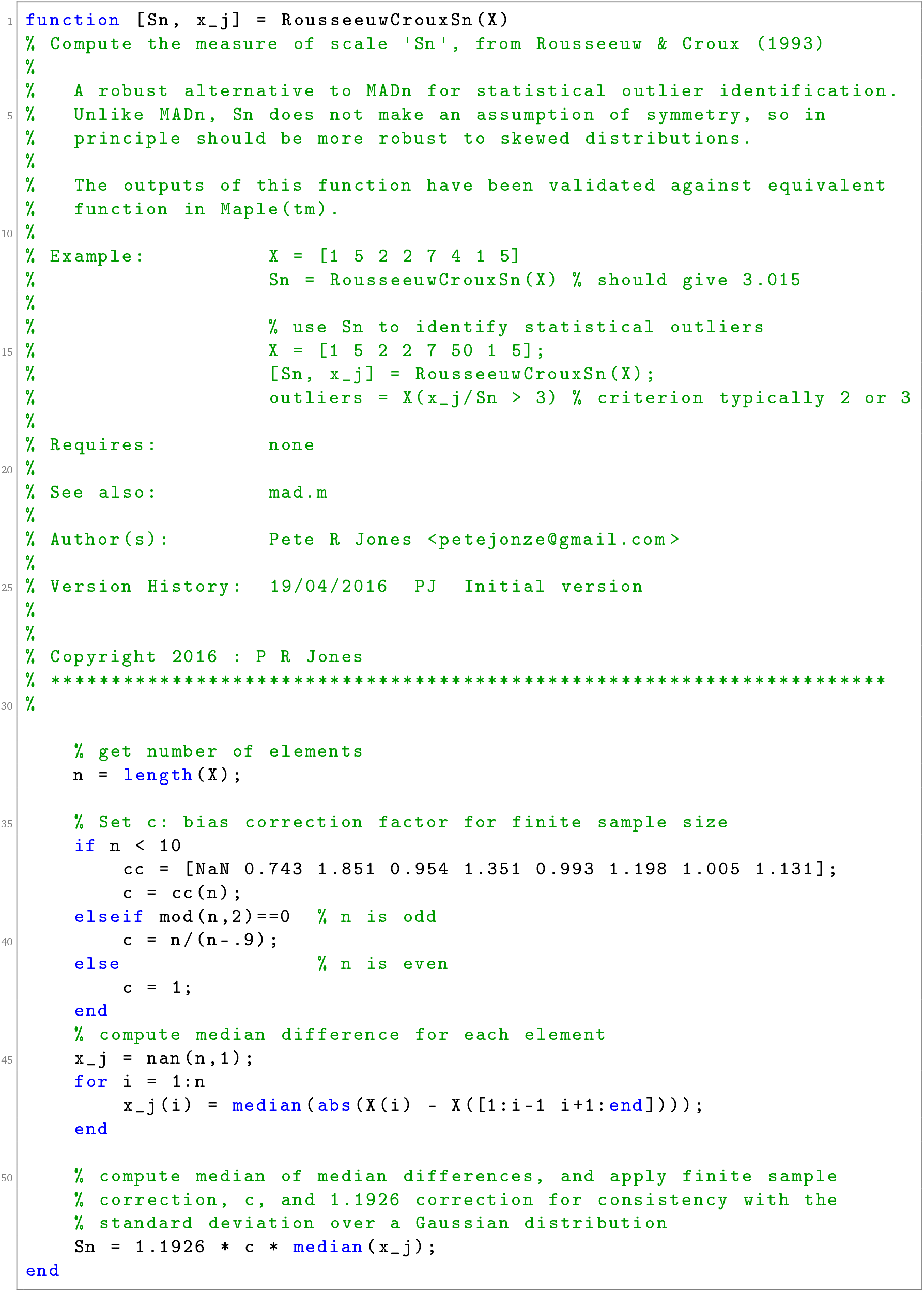
MATLAB code for computing Rousseeuw & Croux’s *S*_*n*_ factor.

### On the ethics of excluding statistical outliers

Excluding outliers is often regarded as poor practice. As shown in *Section 1*, however, the exclusion of outliers can sometimes be preferable to reporting misleading results. Automated methods of statistical outlier identification should never be used blindly though, and they are not a replacement for common sense. Where feasible, datapoints identified as statistical outliers should only be excluded in the presence of independent corroboration (e.g., experimenter observation). Furthermore, best practice dictates that when outliers are excluded, they should continue to be shown graphically, and all statistical analyses should be run twice: with and without outliers included.

## Acknowledgments

This work was supported by the NIHR Biomedical Research Centre located at (both) Moorfields Eye Hospital and the UCL Institute of Ophthalmology.

